# Opposing influence of top-down and bottom-up input on different types of excitatory layer 2/3 neurons in mouse visual cortex

**DOI:** 10.1101/2020.03.25.008607

**Authors:** Rebecca Jordan, Georg B. Keller

**Affiliations:** Friedrich Miescher Institute for Biomedical Research, Basel, Switzerland; Faculty of Natural Sciences, University of Basel, Basel, Switzerland

## Abstract

Processing in cortical circuits is driven by combinations of cortical and subcortical inputs. These signals are often conceptually categorized as bottom-up input, conveying sensory information, and top-down input, conveying contextual information. Using intracellular recordings in mouse visual cortex, we measured neuronal responses to visual input, locomotion, and visuomotor mismatches. We show that layer 2/3 (L2/3) neurons compute a difference between top-down motor-related input and bottom-up visual flow input. Most L2/3 neurons responded to visuomotor mismatch with either hyperpolarization or depolarization, and these two response types were associated with distinct physiological properties. Consistent with a subtraction of bottom-up and top-down input, visual and motor-related inputs had opposing influence in L2/3 neurons. In infragranular neurons, we found no evidence of a difference-computation and responses were consistent with a positive integration of visuomotor inputs. Our results provide evidence that L2/3 functions as a bidirectional comparator of top-down and bottom-up input.

## INTRODUCTION

Learning the relationship between body movements and the resulting sensory feedback is one of the fundamental tasks that the nervous system performs. Predicting the sensory consequences of self-motion is a central component of feedback guided motor control and allows the brain to infer whether sensory stimuli are self-generated or externally generated (Crapse and Sommer, 2008). Exactly how the nervous system learns and represents the relationships between movement and sensory feedback, however, is still unclear. One of the brain structures critically involved in more complex forms of sensorimotor learning is neocortex. A ubiquitous feature of cortical areas is that they receive both bottom-up sensory-driven input, and top-down input that is thought to signal contextual and motor information. The primary visual cortex of mice receives both bottom-up visual input from the lateral geniculate nucleus, as well as top-down input from a variety of other cortical areas, including higher order visual cortices, retrosplenial cortex and anterior cingulate cortex. These top-down inputs have been shown to convey various non-visual types of information, including eye-movement related signals (McFarland et al., 2015), spatial information (Fiser et al., 2016; Saleem et al., 2018), head-direction (Vélez-Fort et al., 2018), and locomotion-related signals (Leinweber et al., 2017). Integrating movement-related input with visual input could allow the cortex to make an accurate inference about how the animal is moving through the world.

There are different ideas about the computational purpose of integrating bottom-up and top-down inputs. One idea is that the top-down input associated with locomotion may function to increase the signal-to-noise ratio of visual responses. This was based on the finding that locomotion results in an increase of visual responses (Niell and Stryker, 2010), as well as a general increase in membrane potential (Bennett et al., 2013; Polack et al., 2013). Neuromodulatory inputs are more active during locomotion (Larsen et al., 2018), with cortex-wide changes in extracellular potassium (Rasmussen et al., 2019), indicating that there is a locomotion-related state change within and across cortical circuits that is likely to underlie this increased gain of visual responses (Fu et al., 2014; Polack et al., 2013). A second idea is that V1 may integrate positively weighted sums of locomotion speed and visual flow speed to estimate an animal’s speed through the world (Saleem et al., 2013). This was based on the finding that neurons in V1 can also be driven in complete absence of visual input in the dark (Keller et al., 2012) and often have activity that correlates positively with both visual flow speed and locomotion speed (Saleem et al., 2013). Another idea is that layer 2/3 (L2/3) neurons use a difference between bottom-up visual input and a top-down prediction to compute visuomotor prediction errors (Keller and Mrsic-Flogel, 2018). This was based on the finding that a subset of L2/3 neurons strongly respond to a sudden mismatch between visual flow feedback and locomotion speed (Keller et al., 2012; Zmarz and Keller, 2016). This last interpretation is at the core of the framework of predictive processing. One of the key features of this model are prediction-error neurons, which receive two sources of input: top-down inputs that convey a prediction of sensory input, and bottom-up inputs that carry sensory-driven information. By comparing these inputs, prediction error neurons are responsive to differences between the two. To do this, top-down and bottom-up inputs could be compared using a divisive mechanism (Spratling, 2008; Spratling et al., 2009), or by a subtractive mechanism (Keller and Mrsic-Flogel, 2018; Rao and Ballard, 1999). A subtractive mechanism would predict that in any given neuron the weights of two types of input are opposing. If bottom up sensory input is excitatory, the top down input should be inhibitory, and vice versa.

The predictive processing framework postulates the existence of two types of prediction error neurons: positive prediction error neurons that subtract a top-down prediction from the sensory input, and negative prediction error neurons that subtract sensory input from the top-down prediction (Keller and Mrsic-Flogel, 2018; Rao and Ballard, 1999). The mismatch-responsive neurons found in L2/3 using calcium imaging are consistent with the latter type of neurons. Visual cortex receives bottom-up excitation and visually driven bottom-up inhibition (Attinger et al., 2017), as well as a diverse combination of excitatory and inhibitory top-down inputs (Gilbert and Li, 2013; Leinweber et al., 2017; Zhang et al., 2014). If the strengths of the two sources of input were balanced and opposing in individual L2/3 excitatory neurons, prediction error responses would arise simply as a result of a temporary imbalance between the two inputs.

To test for the existence of a balance between top-down and bottom-up input in L2/3 and infragranular neurons, we performed intracellular recordings in visual cortex of mice exploring a virtual reality environment. We show that there are two types of responses to visuomotor mismatch in L2/3 excitatory neurons: hyperpolarizing and depolarizing. These two response types are associated with differences in electrophysiological properties, visual responses, and membrane potential dynamics during locomotion indicating they may be different neuron types. Moreover, the sign of the response to visual input and the sign of the response to locomotion-related input was inversely related, consistent with L2/3 computing a difference between the two inputs. By contrast, deep layer neurons did not show this characteristic, instead exhibiting positive signs of responses to both types of input. Thus, we demonstrate layer and neuron-type specific integration of sensory and motor-related inputs.

## RESULTS

To assess subthreshold responses to visuomotor mismatch, we made blind whole cell recordings in visual cortex of awake mice, head-fixed on a spherical treadmill with locomotion coupled to visual flow feedback in a virtual reality environment (**Figure 1A**) (Leinweber et al., 2014). To ensure sufficient levels of locomotion during recording experiments, mice were habituated to this paradigm in several sessions prior to whole cell recordings (**Figure 1B**). During each whole cell recording experiment, the mouse first experienced full-field visual flow that was coupled to its locomotion. The coupling of visual flow to locomotion was interrupted by suddenly halting visual flow for 1 s at random times to generate visuomotor mismatch events (Keller et al., 2012). Subsequently, we decoupled locomotion and visual stimuli to measure the independent contributions of locomotion and visual flow on neuronal activity. Visual stimuli consisted of full-field, fixed-speed flow of the virtual tunnel walls, presented at random times for 1s, regardless of the mouse’s locomotion behavior. We made whole cell recordings from a total of 54 neurons. We focused our analysis on putative excitatory neuron types by excluding neurons with high input resistance (greater than 100 MΩ), as well as neurons with a spike half-width below 0.6 ms (Gentet et al., 2012; Pala and Petersen, 2015). By doing this, we excluded 15% of neurons in total, leaving 46 putative excitatory neurons in the sample (**Figure S1A-C**). We then restricted the dataset to 32 putative L2/3 excitatory neurons by analyzing only neurons recorded at less than 400 µm vertical depth from the brain surface.

**Figure 1.**
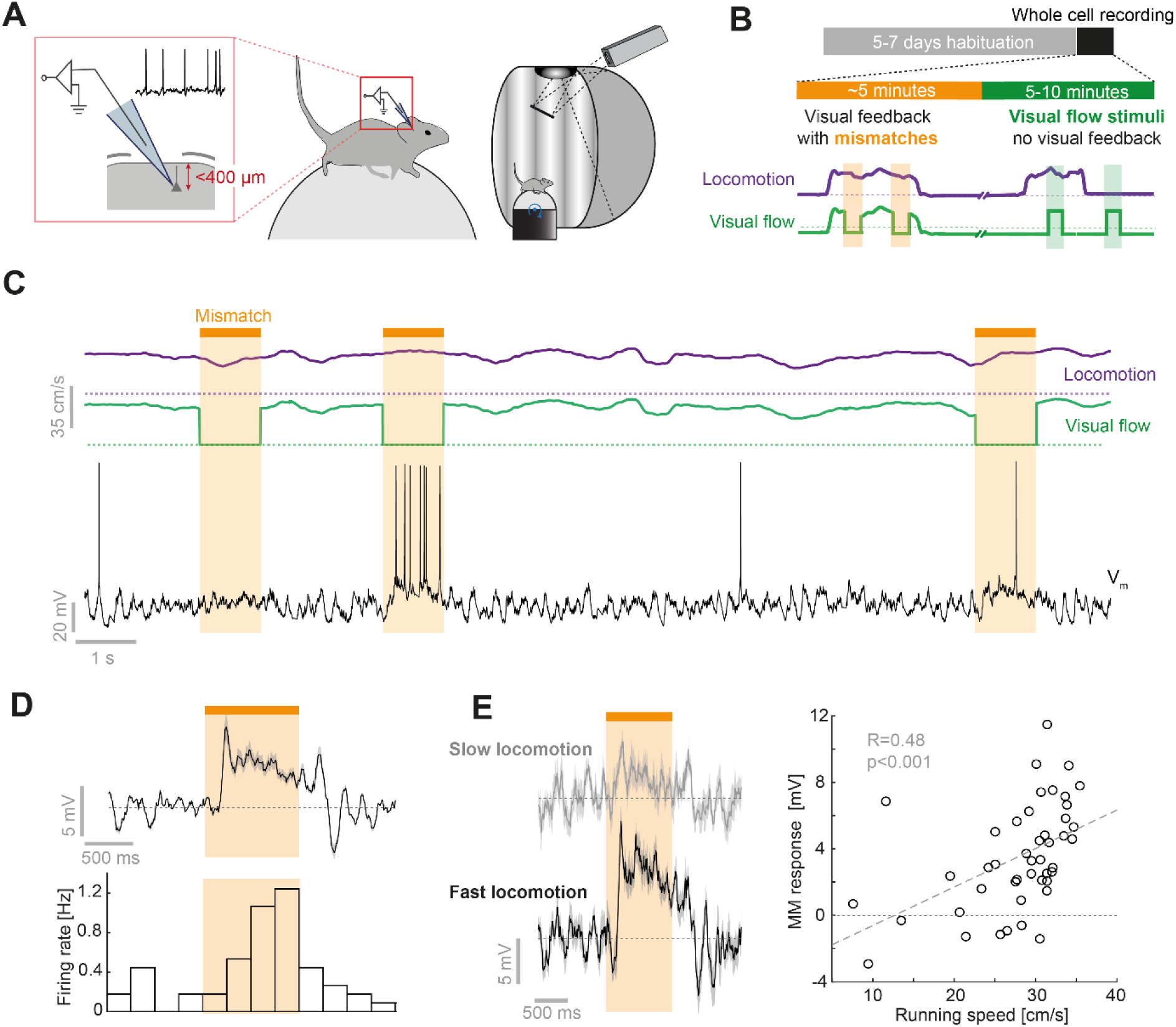
Whole cell recordings during visuomotor coupling and mismatch. (**A**) Schematic of whole cell recordings at L2/3 depth and of the visuomotor virtual reality set up. (**B**) Mice were habituated on the setup for 5 to 7 days before whole cell recording experiments. Experiments consisted of an initial closed-loop phase, during which visual flow feedback was coupled to the mouse’s locomotion interspersed with sudden unpredictable visual flow halts (mismatches). In a second phase, visual flow was presented in 1 s pulses of visual flow independent of locomotion. (**C**) Top: Locomotion (purple) and visual flow (green) speeds during a visuomotor closed-loop session. Dashed lines mark zero speed. Visuomotor mismatch events are marked by an orange bar and shading. Bottom: Example membrane potential trace from a depolarizing mismatch (dMM) neuron. (**D**) Average membrane potential response (top) and firing rate (bottom) histogram for mismatch trials for example neuron shown in **C** (average over 45 mismatch events). Shading indicates SEM. (**E**) Left: Average membrane potential response for the 10 mismatch events with highest locomotion speed (black) and the 10 mismatch events with lowest locomotion speed (gray), for the example neuron shown in **C**. Shading indicates SEM. Right: Scatter plot between average locomotion speed in 2 s prior to mismatch and membrane potential response to mismatch for 45 trials recorded in example neuron shown in **C**. Dashed gray line shows a linear regression.

### Subthreshold mismatch responses are widespread in putative L2/3 excitatory neurons and distinguish different neuron types

We first assessed the responses of putative L2/3 excitatory neurons to visuomotor mismatch (15 ± 10 (mean ± standard deviation) mismatch presentations per neuron). Consistent with mismatch responses described previously using calcium imaging, we found neurons with clear depolarizing responses to visuomotor mismatch stimuli (**Figure 1C and 1D**). The strengths of these responses were often correlated with the running speed of the mouse immediately preceding the mismatch event (**Figures 1E- and 1F, S1D and S1E**), with the depolarization becoming stronger with faster locomotion speeds. This is congruent with the responses reflecting the degree of error between running speed and visual flow speed, and consistent with the responses found using calcium imaging (Zmarz and Keller, 2016).

In total, 17 of 32 neurons significantly responded with at least 1 mV average depolarization and 6 of 32 neurons responded with at least 1 mV hyperpolarization to visuomotor mismatch (**Figures 2A-C**). We will refer to these neurons as depolarizing mismatch (dMM) neurons and hyperpolarizing mismatch (hMM) neurons respectively. The remaining neurons (9 of 32) did not exceed the average 1 mV threshold but were not unresponsive and often showed brief depolarizing responses at the onset and/or the offset of mismatch. Spiking responses were less common, with only 3 of 32 neurons displaying more than 0.25 spikes per mismatch stimulus on average (**Figure 2D**). Supporting the idea that hMM and dMM neurons are different neuron types driven by different input pathways, we found that electrophysiological properties differed between dMM and hMM neurons. dMM neurons had more depolarized resting membrane potentials (mean ± SD, dMM: -77.7 ± 7.2 mV, hMM: -90.4 ± 3.7 mV, p < 0.01, Student’s t-test; **Figure 2E**), more depolarized spike thresholds (mean ± SD, dMM: -34.7 ± 4.4 mV, hMM: -41.1 ± 2.1 mV; p < 0.05, Student’s t-test; **Figure 2F**) and lower spike rates than hMM neurons (dMM: median spike count = 0, IQR = 0 to 0.0008 spikes; hMM: median spike count = 0.05, inter-quartile range (IQR) = 0 to 0.11 spikes; p < 10^−7^, Brown-Forsythe test; p < 0.03, Wilcoxon rank sum test; **Figure 2G**). We found no significant differences in input resistances or membrane time constants between the two groups (**Figure S2**), however, using multiple linear regression involving five variables (input resistance, membrane time constant, resting membrane potential, baseline firing rate and membrane potential variance when stationary), we could explain almost 70% of the variance in mismatch responses (R^2^ = 0.68, p < 0.001, 23 neurons). Thus, most L2/3 neurons responded to visuomotor mismatches, and the sign of this response to mismatch correlates with differences in electrophysiological properties.

**Figure 2.**
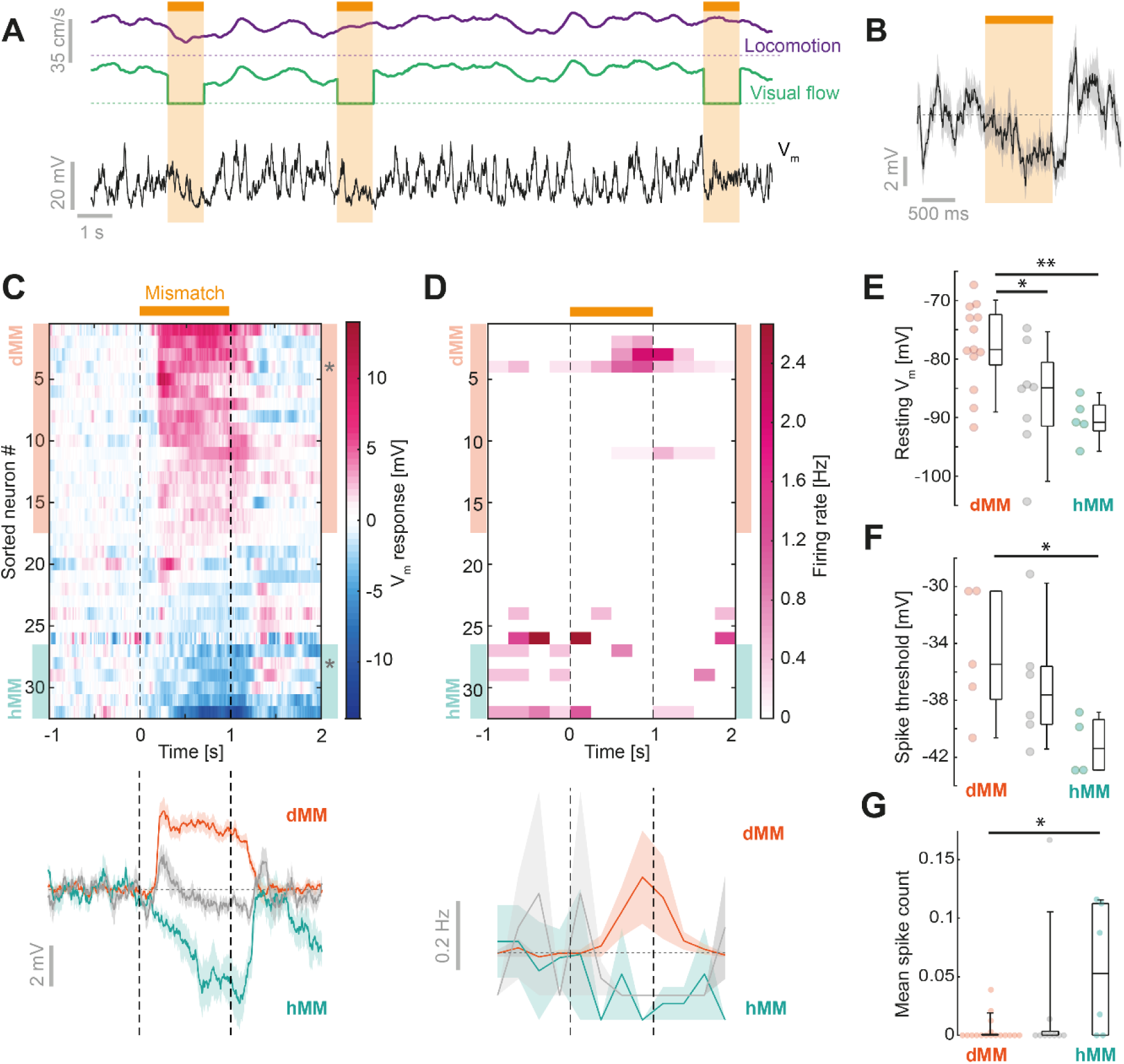
Subthreshold mismatch responses are widespread in L2/3 and the direction of response correlates with electrophysiological properties. (**A**) Locomotion (purple), visual flow (green) and membrane potential (black) traces from a neuron with a hyperpolarizing response to mismatch. Dotted lines indicate 0 cm/s for visual flow (green) and locomotion (purple). Mismatch events are marked by an orange bar and shading. (**B**) Average membrane potential response to mismatch for the example neuron in **A**. Shading indicates SEM over 15 trials. (**C**) Top: Heatmap of average membrane potential (V_m_) responses to mismatch from all L2/3 neurons. Neurons are sorted by average mismatch response. Neurons classified as depolarizing mismatch (dMM) and hyperpolarizing mismatch (hMM) are marked by orange and turquoise shading respectively. Bottom: Average response across 17 dMM neurons (orange), 6 hMM neurons (turquoise), and the remaining 9 unclassified neurons (gray). Asterisks indicate example neurons shown in panel **A**, and **Figure 1A**. (**D**) Top: Heatmap of average firing rate during mismatch. Color coding and sorting as in **C**. Bottom: Average mismatch induced change spike count for the same groups of neurons as in **C**. (**E**) Resting membrane potential recorded just after entering whole cell recording mode for 5 hMM neurons, 13 dMM neurons, and 8 unclassified neurons. *: p < 0.05, **: p < 0.01, Student’s t-test. Box plots show median, quartiles, and range excluding outliers. (**F**) As in **E**, but for spike threshold. Note, only neurons with spontaneous spikes were included. *: p < 0.05, Student’s t-test. (**G**) As in **E**, but for baseline spike count in a window 2 s prior to mismatch onset. *: p < 0.05, Wilcoxon rank sum test.

### Mismatch responses are anticorrelated with visual flow responses in putative L2/3 excitatory neurons

The observed mismatch responses either could be computed in L2/3 neurons or could be inherited from other layers in visual cortex, or from other brain regions that provide input to L2/3 visual cortex neurons. If they are computed locally and arise from a reduction in bottom-up visually driven input, we would expect to see an opposing relationship between the sign of the mismatch response and that of the response to visual flow. If the bottom-up visual input is depolarizing, mismatch responses should be hyperpolarizing, and vice versa. To examine this, we analyzed the responses of all neurons to visual input in the form of brief periods of visual flow, presented independently from locomotion. Since the visual flow responses during locomotion and during stationary periods were correlated (**Figure S3**), we included all presentations in our analysis in order to maximize trial number. Of the 32 putative L2/3 we lost 5 neurons before being able to record visual responses. Consistent with an opposing response to mismatch and visual input, we found that on average dMM neurons responded to these visual flow stimuli with a hyperpolarization (**Figure 3A-C**), while hMM neurons responded to visual flow stimuli with depolarization (**Figure 3C**). Of 27 neurons recorded, only 3 exhibited spiking responses to visual stimuli and were all hMM neurons. The subthreshold response difference to visual input was significantly different in dMM and hMM neurons (mean ± SD, dMM: -0.26 ± 1.57 mV, n = 14; hMM: 3.42 ± 2.91 mV, n = 5; p < 0.003, Student’s t-test; **Figure 3D**). Across all neurons, we found a significant negative correlation between visual flow response and mismatch response (R = -0.50, p < 0.01, n = 27, **Figure 3E**).

**Figure 3.**
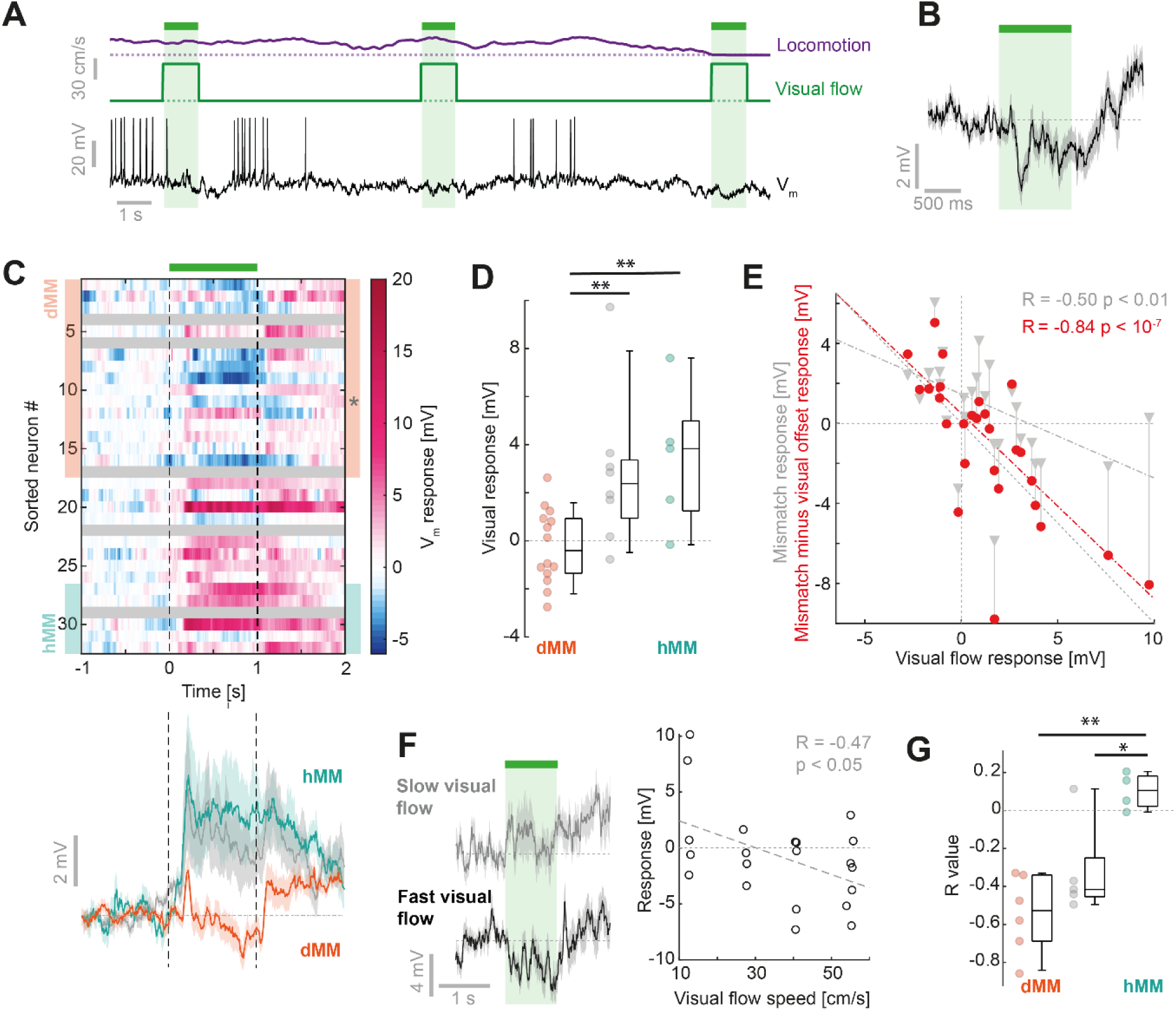
Visual flow responses inversely relate to mismatch responses in L2/3 neurons. (**A**) Example membrane potential trace (black) recorded from a dMM neuron during visual flow presentations. Brief 1 s visual flow stimuli (shaded green) evoked hyperpolarization. Locomotion is shown in purple. (**B**) Average membrane potential response to visual flow stimuli for the example neuron in **A**. Gray shading indicates SEM over 42 trials. (**C**) Top: Heatmap of average membrane potential response to full-field visual flow stimuli across of 27 L2/3 neurons. Responses are sorted according to mismatch response, as in **Figure 2C**. Gray shading marks neurons for which we did not have sufficient data in the open-loop condition. Bottom: Average response across 17 dMM neurons (orange), 6 hMM neurons (turquoise), and the remaining 9 neurons (gray). Asterisk indicates example neuron shown in panel **A**. (**D**) Average V_m_ responses to 1 s visual flow, compared for 5 hMM, 14 dMM, and 8 unclassified neurons. **: p < 0.01, Student’s t-test. (**E**) Scatter plot between average visual response and average mismatch response (gray triangles) for 27 neurons. For red data points, mismatch responses are corrected for visual flow offset responses by subtracting the average response to visual flow offset from the mismatch response. Dashed gray and red lines are linear fits to the respective data. (**F**) Left: Average response to visual flow stimuli of lowest two visual flow speeds (gray, 7 trials), and highest two speeds (black, 11 trials) for an example neuron. Shading indicates SEM over trials. Right: Scatter plot between visual flow speed and membrane potential response to visual flow for 18 trials of the example neuron. (**G**) Correlation coefficients between visual flow speed and membrane potential response compared for 6 hMM neurons, 4 dMM neurons and 5 unclassified neurons. *: p < 0.05, **: p < 0.01, Student’s t-test.

In addition to responses to visual flow onset, we found that many neurons also exhibited depolarizing responses or persisting depolarization after visual flow offset (**Figure 3C**). One interpretation of this is that the offset of visual flow results in separate excitatory input to L2/3 neurons. Consequently, mismatch responses would be a combination of the reduction of a bottom-up input correlated with visual flow and a parallel increase in input driven by the visual flow reduction. Correcting the mismatch responses for this visual flow offset response revealed an almost perfect balance between opposing visually driven responses and mismatch responses across the population of L2/3 neurons (R = -0.84, p < 10^−7^, n = 27; **Figure 3E**; see Methods).

In a subset of neurons, we tested the effect of varying the visual flow speed and presented four distinct visual flow speeds (**Figure 3F**). For dMM neurons, visually driven hyperpolarization scaled with visual flow speed, resulting in negative correlations between visual flow speed and membrane potential response (mean R value ± SD, = -0.54 ± 0.21, n = 6) (**Figure 3G**). By contrast, in hMM neurons the correlation of visually driven depolarization with visual flow speed was positive, consistent with an excitatory bottom-up input correlated with visual flow speed (mean R value ± SD = 0.1 ± 0.1, n = 4; dMM vs hMM: p < 0.003, Student’s t-test). This balance between opposing visual and mismatch responses is consistent with mismatch responses arising from transient removal of bottom-up visual input.

### The influence of locomotion on membrane potential differs depending on mismatch response

Computing visuomotor prediction errors requires a top-down input to convey a prediction of visual flow given movement. A potential source for such a top-down input to V1 is A24b/M2 (Leinweber et al., 2017). If this were the case, we would expect to find a motor-related input to a given neuron whose strength is correlated with the strength of the mismatch responses and anti-correlated with the strength of visual response. Thus, dMM neurons should receive motor-related excitation, while hMM neurons should receive motor-related inhibition (Keller and Mrsic-Flogel, 2018). Complicating this is the fact that locomotion is associated with a brain state change, likely driven by neuromodulatory inputs (Fu et al., 2014; Polack et al., 2013). We first analyzed the membrane potential changes driven by locomotion and, consistent with previous work, found a systematic change in both mean membrane potential (V_m_) and variance (**Figures 4A-C and S4**) (Bennett et al., 2013; Polack et al., 2013). The membrane potential became more depolarized (mean ± SD, ΔV_m_= 4.5 ± 2.5 mV, p < 10^−10^, 39 neurons, paired t-test) and less variable (mean ± SD, ΔV_m_ SD= -1.8 ± 1.5 mV, p < 10^−8^, paired t-test) during locomotion, with only a small change in spike rates (mean ± SD, ΔFR = 0.11 ± 0.65 Hz, p = 0.31, paired t-test). Quantifying the membrane potential changes driven by locomotion onset in absence of coupled visual flow in the open loop condition, we found that all neurons displayed locomotion-related depolarization of membrane potential that began prior to locomotion onset (**Figure 4C**). This is consistent with locomotion onset responses observed in suprathreshold responses in V1 (Keller et al., 2012; Saleem et al., 2013), and is likely caused by neuromodulatory input (Polack et al., 2013). We also found that locomotion caused visual responses to become significantly more depolarizing (**Figure S3**), consistent with previous findings (Bennett et al., 2013). Although both dMM and hMM neurons both depolarized during locomotion, locomotion onset responses correlated positively with mismatch responses (**Figure 4E**). These results are consistent with a neuromodulatory state change causing widespread depolarization of neurons, alongside a separate locomotion-related drive that correlates with mismatch response. To better separate the effects of the state change from a potential direct locomotion-related drive, we analyzed the correlation between membrane potential and locomotion speed of the mouse only during times of locomotion, to minimize the influence of state transitions associated with locomotion on- and offsets (see Methods; **Figure S6**). When comparing responses in hMM and dMM neurons restricted to times of locomotion, we found that dMM neurons depolarized with increasing locomotion speed, while hMM neurons hyperpolarized with increasing locomotion speed (dMM: mean R value ± SD = 0.14 ± 0.13, n = 12; hMM: -0.06 ± 0.07, n = 5; p < 0.005, Student’s t-test; **Figure 4E**). This would be consistent with a locomotion-related excitation onto dMM neurons and a locomotion-related inhibition onto hMM neurons that both scale with locomotion speed, in addition to the state-dependent depolarization. Lastly, if mismatch responses are the result of opposing visual flow and locomotion speed inputs, the correlation of the membrane potential with locomotion and that of membrane potential with visual flow speed should have opposite sign. This was indeed the case, the membrane potential of dMM neurons exhibited negative correlations with visual flow and positive correlations with locomotion speed, and the opposite was true in hMM neurons (**Figures 4F and S6**). Consistent with a balance of two opposing inputs, we found that in both hMM and dMM neurons the timing of the peak cross-correlation of membrane potential with locomotion and was well matched with that of the cross-correlation of membrane potential with visual flow. These data are consistent with L2/3 mismatch-responsive neurons computing a comparison between a visual flow input and a locomotion related input using balanced and opposing excitatory and inhibitory input.

**Figure 4.**
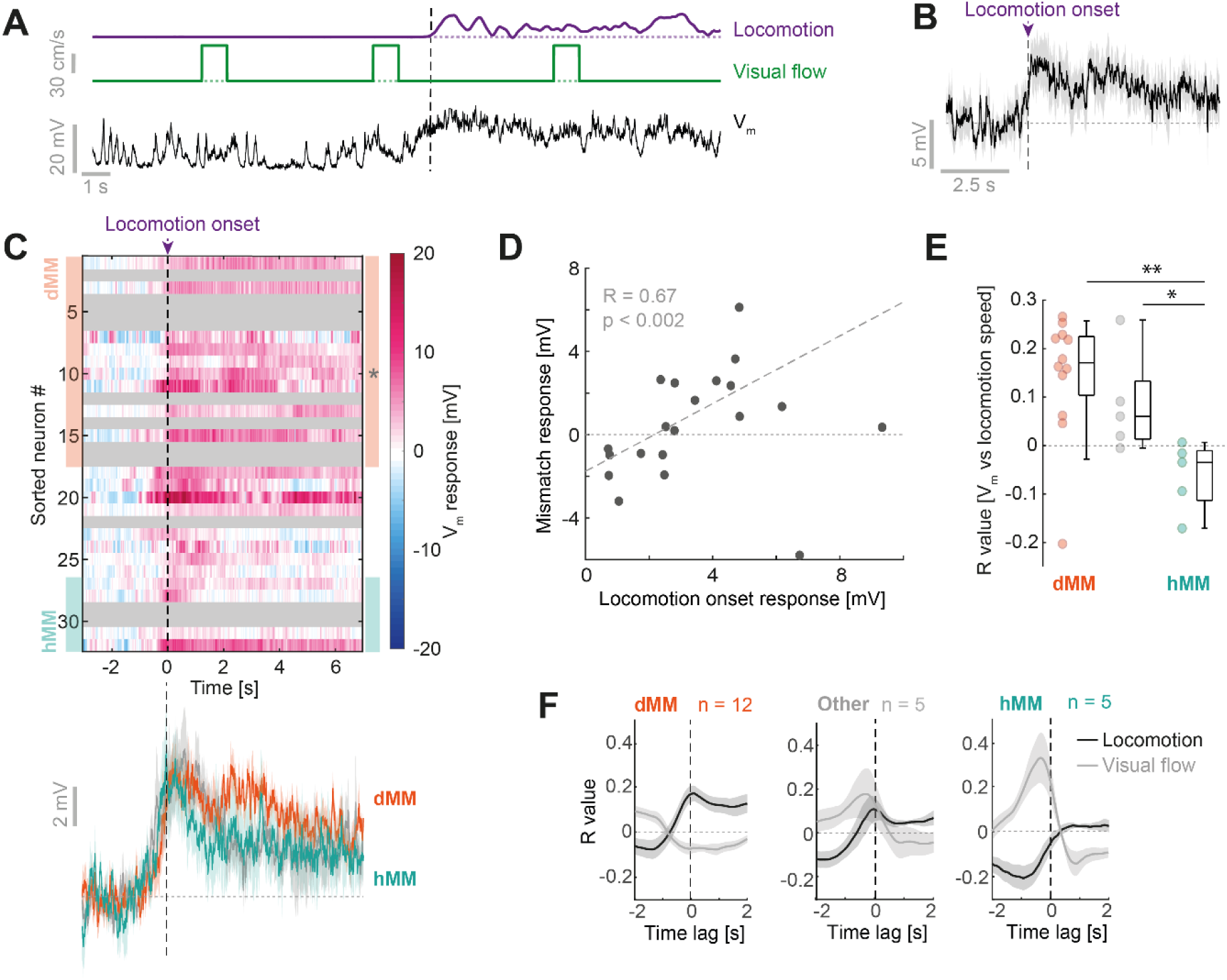
The influence of locomotion on membrane potential differs depending on mismatch response. (**A**) Example membrane potential trace (black) from a neuron during locomotion onset (vertical dashed line) in the open-loop condition. (**B**) Average V_m_ response to locomotion onset (purple dashed line) for example neuron in **A**. Shading indicates SEM over 4 trials. (**C**) Top: Heatmap of average membrane potential responses to locomotion onset for all L2/3 neurons. Responses are sorted as in **Figure 2C**, according to the average mismatch response. Gray shading marks neurons for which we did not have sufficient data in the open-loop condition. Bottom: Average response across 9 depolarizing mismatch neurons (orange), 4 hyperpolarizing MM neurons (turquoise) and the 8 remaining neurons (gray). Shading indicates SEM over neurons. (**D**) Average locomotion onset response (0 to 6 s after locomotion onset) plotted against mismatch response for 21 neurons. (**E**) Correlation coefficients between locomotion speed and V_m_ compared for 12 dMM neurons, 5 hMM neurons and remaining 5 neurons. **: p < 0.01, Student’s t-test. (**F**) Average cross correlations between membrane potential and locomotion speed (black) or visual flow speed (gray). Negative time values indicate locomotion/visual speed preceding membrane potential. Shading indicates SEM over neurons.

### Infragranular layers positively integrate visual and motor-related input differently from L2/3

Based on physiological and anatomical characteristics, it has been suggested that infragranular layers 5 and 6 (L5/6) neurons perform a computational function different from L2/3 neurons. L5 neurons, for example, are strongly and broadly interconnected and appear to employ a dense firing code, while L2/3 are more weakly interconnected, fire sparsely and exhibit more prominent lateral inhibition (Harris and Mrsic-Flogel, 2013). To examine whether mismatch responses are computed in a layer-specific fashion, we compared the L2/3 responses to those of neurons recorded at a depth of between 480 and 750 µm from the brain surface (**Figure 5A**), which we will consider putative L5/6 neurons (n = 14). The electrophysiological properties of these neurons differed from those of L2/3 neurons: L5/6 neurons had significantly higher input resistances, lower spike thresholds, and tended to spike more than L2/3 neurons (**Figure S5**). These differences are consistent with those previously found between L5 and L2/3 in neocortex (De Kock and Sakmann, 2009; Lefort et al., 2009; Sakata and Harris, 2009). We next analyzed the responses of L5/6 neurons to visuomotor mismatch (**Figures 5B and 5C**). In stark contrast to L2/3 neurons, depolarizing responses to mismatch were rare in L5/6 (2 of 14 neurons), while half of the neurons (7 of 14) showed a hyperpolarizing response (**Figure 5D**). This resulted in a significant difference between average mismatch responses in L2/3 and L5/6 (mean ± SD, L2/3: 0.8 ± 2.4 mV, 32 neurons; L5/6: -1.3 ± 2.2 mV, 14 neurons; p < 0.01; Student’s t-test; **Figures 5E and 5F**). This paucity of depolarizing mismatch responses in L5/6 could be the result of a) a reduced bottom-up visual inhibition, b) reduced excitatory locomotion-related input, c) a lack of balanced and opposing tuning between these two sources of input, or d) any combination of the above. To examine these possibilities, we analyzed visual flow responses and found that most L5/6 neurons exhibited depolarizing responses to visual flow stimuli (**Figure 5G**). The distribution of average visual responses did not significantly differ from that of L2/3 neurons (mean ± SD, L2/3: 1.3 ± 2.8 mV, 27 neurons; L5/6: 2.0 ± 1.9 mV, 13 neurons; p = 0.47, Student’s t-test; **Figures 5H and 5I**), although hyperpolarizing visual responses did not occur in L5/6 neurons (L2/3: 26%, L5/6: 0%). Next, we looked at locomotion onset responses in L5/6 neurons. As with L2/3 neurons, nearly all L5/6 neurons underwent a depolarization at locomotion onset (**Figures 5J and 5K**) and displayed similar changes in membrane potential dynamics (**Figure S4**). However, on average this depolarization was smaller in L5/6 neurons than that of L2/3 neurons (mean ± SD, L2/3: 3.6 ± 2.3 mV; L5/6: 2.0 ± 2.7 mV; p = 0.09, Student’s t-test; **Figure 5L**). We also found that locomotion had distinct effects on the visual response in L5/6 versus L2/3 neurons: while locomotion caused visual responses to be more depolarizing in L2/3 neurons, there was no consistent effect in L5/6 neurons (**Figure S3**), similar to findings in somatosensory cortex (Ayaz et al., 2019). Finally, we compared how the membrane potential in L5/6 neurons scaled with locomotion speed and visual flow speed. Unlike for L2/3 neurons, the membrane potential correlated positively with visual flow speed and locomotion speed for most L5/6 neurons (10 of 12 **Figures 6A and S5**). Plotting the correlation of membrane potential with visual flow against that of membrane potential with locomotion speed for each neuron, we found a significant anticorrelation between these values in the L2/3 population (R = -0.67, p < 0.001, 22 neurons; **Figure 6A**). In the L5/6 population we found no significant correlation, with most neurons in the top right quadrant of positive correlation with both visual flow speed and locomotion speed (R = -0.26, p = 0.42, 12 neurons). Representing each neuron as an angle based on the two correlation values, we found that L2/3 showed a significantly higher proportion of neurons with angles corresponding to opposing signs of correlation (90-180 degrees and 270-360 degrees) than L5/6 neurons (L2/3: 17 of 22 neurons; L5/6: 2 of 12 neurons; p=0.001, Fisher’s exact test; **Figure 6B**). The distribution of these angles for L2/3 and L5/6 neurons were significantly more anticorrelated than expected by chance (p < 0.02, see Methods). Across neurons, the difference between the correlation of membrane potential with locomotion and the correlation of membrane potential with visual flow was a good predictor of mismatch responses in L2/3 (R = 0.59, p < 0.01), but not in L5/6 (R = -0.06, p = 0.84) (**Figure 6C**). Thus, the absence of depolarizing mismatch responses in L5/6 neurons is likely a result of a combination of a reduced bottom-up inhibition, and a lack of a balanced and opposing tuning between visual and locomotion related inputs. In sum, widespread, strong mismatch responses and the opposing visual and motor-related inputs necessary to compute them are a specific feature of L2/3 and are absent in L5/6 neurons, indicative of separate computational roles of L2/3 and L5/6 in visuomotor integration.

**Figure 5.**
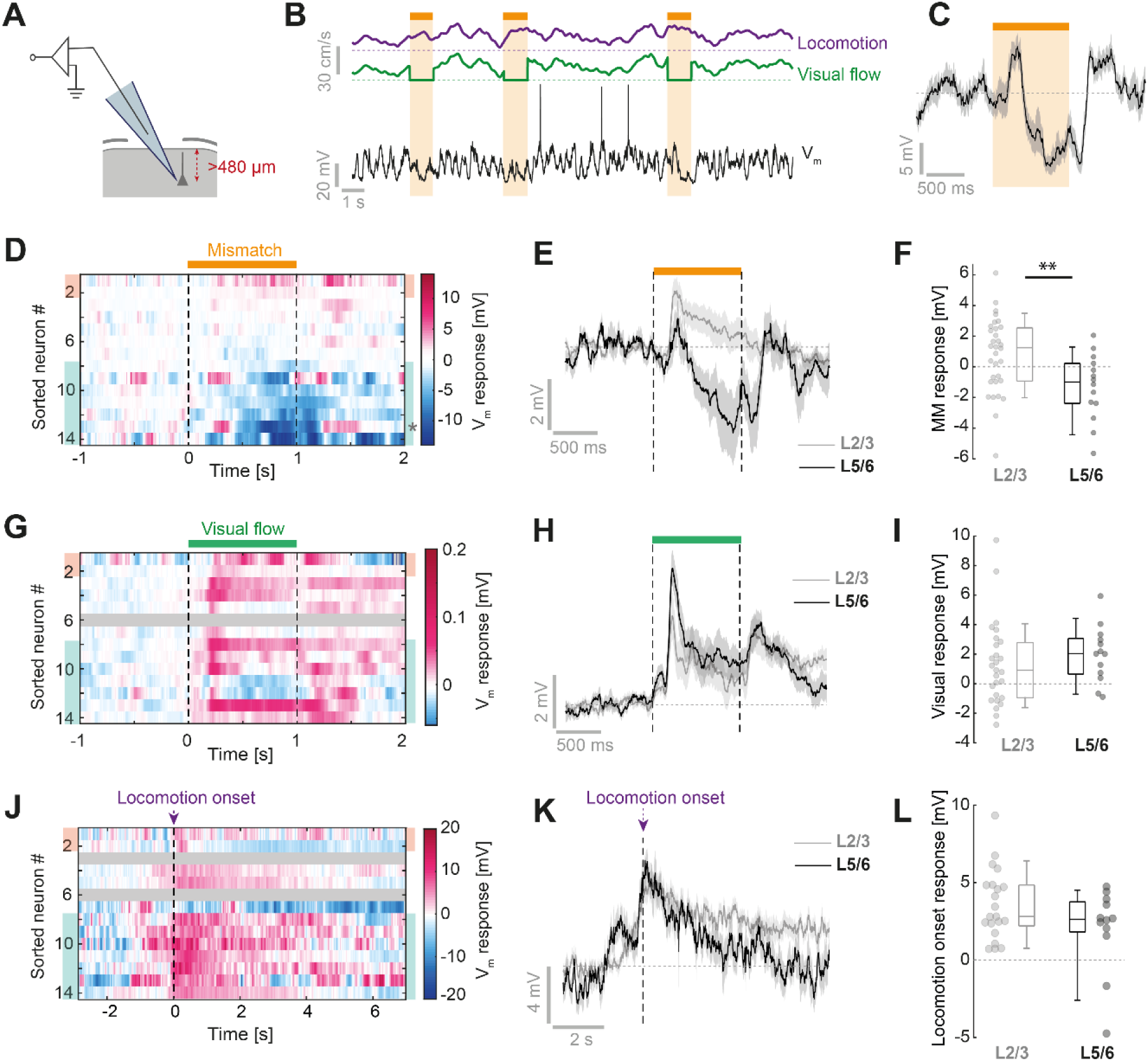
Deep layer neurons show lack of depolarizing mismatch responses and visuomotor integration differs from L2/3 neurons. (**A**) Neurons recorded at a vertical depth greater than 480 µm were classified as putative L5/6 neurons. (**B**) Example membrane potential trace from a putative L5/6 neuron recorded during visuomotor coupling with mismatch stimuli (orange bar and shading). (**C**) Average membrane potential response to mismatch stimulus for the neuron in **B**. Shading indicates SEM over 13 trials. (**D**) Heatmap of average membrane potential responses to mismatch of all L5/6 neurons. Neurons are sorted by average mismatch response. (**E**) Average response to mismatch across all L5/6 neurons (black, 14 neurons), compared to the average response to mismatch across all L2/3 neurons (gray, 32 neurons). Shading indicates SEM over neurons. (**F**) Average mismatch responses of L2/3 and L5/6 neurons. **: p < 0.01, Student’s t-test. (**G**) As in **D**, but for visual flow responses. (**H**) As in **E**, but for visual flow responses. (**I**) As in **F**, but for visual flow responses. (**J**) As in **D**, but for locomotion onset responses. (**K**) As in **E**, but for locomotion onset responses. (**L**) As in **F**, but for locomotion onset responses.

**Figure 6.**
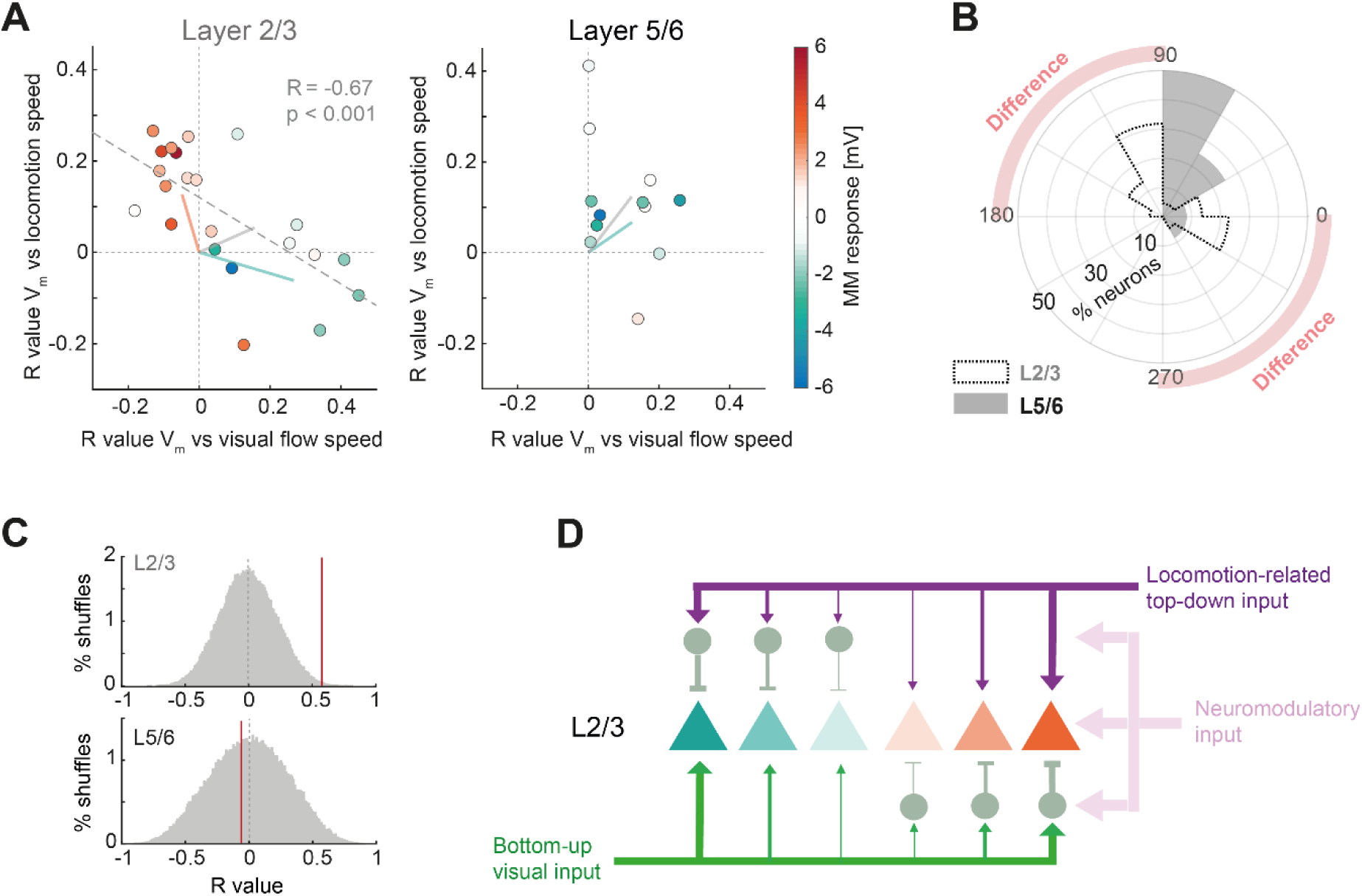
L2/3 neurons, but not L5/6 neurons, integrate locomotion and visual inputs with opposing sign. (**A**) Left: Scatter plot of the correlation coefficient between V_m_ and visual flow speed, and the correlation coefficient between V_m_ and locomotion speed for 22 L2/3 neurons. Right: The same for 12 L5/6 neurons. Data points are colored by mismatch response. Gray dashed line is a linear regression to the data. Pale solid lines indicate average angles for hMM neurons (turquoise), dMM neurons (orange), and remaining neurons (gray). (**B**) Histogram of the angles in **A** for each neuron. The two histograms are significantly anticorrelated compared to shuffled datasets (p < 0.02, see Methods). (**C**) The difference between the correlation of membrane potential with visual flow and the correlation of membrane potential with locomotion speed is a good predictor of mismatch responses in L2/3 neurons (R = 0.59, vertical red line), but not in L5/6 (R = -0.06, vertical red line). The gray histograms are shuffle controls in which the locomotion and visual flow correlation values are scrambled across neurons. (**D**) Schematic of hypothesized L2/3 circuit. Excitatory neurons (triangles) in L2/3 have a range of responses to visuomotor mismatch from strong depolarization (orange) to strong hyperpolarization (green). The strength of this response reflects the balance between feedforward and top-down excitation and inhibition for this particular combination of visuomotor inputs. Locomotion causes both direct excitatory input and disynaptic feed-forward inhibition (via inhibitory interneurons, in gray), as well as a state change that affects neurons via neuromodulatory input. Width of arrows indicates relative strength of input.

## DISCUSSION

A canonical feature of cortical circuits is the pattern of long-range inputs that is conceptually often divided into two types of input: bottom-up and top-down. Bottom-up input originates in thalamus and areas thought to be at a lower level in a local hierarchy of cortical areas and generally conveys sensory information from the periphery. Top-down input originates from areas at a higher level of a local cortical hierarchy and is thought to provide contextual information and underlie phenomena such as selective attention (Busse et al., 2017; Engel et al., 2001; Makino and Komiyama, 2015; Zhang et al., 2014). In some computational frameworks, top-down inputs are thought to convey a prior or prediction of bottom-up input. In the predictive processing framework, top-down predictions are thought to be subtracted from bottom-up input to compute bidirectional prediction errors that are used to update an internal representation (Keller and Mrsic-Flogel, 2018; Rao and Ballard, 1999). By probing intracellular voltage responses to visuomotor mismatches we found that putative L2/3 excitatory neurons fall into different response classes consistent with either bottom-up or top-down excitatory drive. These neurons exhibit a visual flow input of matching strength and opposite sign of motor-related input, consistent with a subtractive prediction-error computation in L2/3. This was not a feature of L5/6 neurons, which largely displayed hyperpolarizing responses to mismatch and appeared to integrate positive valued locomotion-related and visual inputs. These results are consistent with a model of visual cortex in which L2/3 generates sensorimotor prediction errors, and L5/6 integrates L2/3 input to update an internal representation of the world (Keller and Mrsic-Flogel, 2018; Rao and Ballard, 1999).

A classic example of prediction errors can be found in the dopaminergic reward system. Here, reward prediction errors are thought to be encoded bidirectionally in individual neurons with increases and decreases in firing rate corresponding to a positive or negative prediction error (Schultz et al., 1997). In our recordings, at least two kinds of mismatch response are distinguishable in L2/3: those that hyperpolarize and those that depolarize during mismatch. These neurons also show distinct visual responses, relationships between membrane potential and locomotion speed, and electrophysiological properties. These two neuron types could correspond to positive and negative prediction error neurons, which signal that the sensory input is higher or lower than predicted, respectively. Splitting prediction error responses into two separate populations of neurons is necessary when baseline firing rates are low, as they are in L2/3 neurons.

What fraction of neurons that respond to mismatch would we expect to find assuming the predictive processing framework were a useful model? A simple upper bound on this would be at most half of the prediction error neuron population, half responds to positive prediction errors, the other half to negative prediction errors. However, this assumes that we are probing the entire space of all visual prediction errors represented in V1. In our experiments, we probe prediction errors by breaking the coupling between forward locomotion and backward visual flow. This represents only a small fraction of total space of visuomotor coupling a mouse will experience. V1 receives top-down input that conveys predictions of visual input given locomotion (Leinweber et al., 2017), spatial location (Fiser et al., 2016), and visual surround (Keller et al., 2020). In addition, V1 receives inputs that convey vestibular signals (Vélez-Fort et al., 2018) and auditory signals (Ibrahim et al., 2016; Iurilli et al., 2012). In principle, all of these top-down inputs could be associated with a population of prediction error neurons selective for the particular type of error. Suggestive of this is the finding that L2/3 neurons that respond to the omission of a visual input a mouse expects to see based on spatial location are different from those that respond to mismatch (an absence of visual flow expected based on forward locomotion) (Fiser et al., 2016). Thus, one would not expect all prediction error neurons to respond to a single type of prediction error. Using extracellular recordings across the entire cortical depth likely dominated by infragranular responses, a previous study found that only 5 of 73 neurons were selective for the difference between visual speed and locomotion speed (Saleem et al., 2013). The fraction of L2/3 neurons estimated to respond to visuomotor mismatch events based on calcium imaging is in the range of 20% to 30% (Attinger et al., 2017; Keller et al., 2012).

Each neuron likely has some tuning that relates to the top-down input distribution conveying the prediction used to compute the prediction error. This tuning would be reflected in a graded subthreshold response to one particular type of prediction error as a result of balanced bottom-up and top-down input, independent of whether the neuron has a spiking response to that particular prediction error (**Figure 6D**). Consequently, one would expect to find that negative prediction error neurons are net bottom-up inhibited and top-down excited, and vice versa for positive prediction error neurons. We do indeed find that about half of the L2/3 neurons exhibit depolarizing mismatch responses consistent with a negative prediction error input balance. The question remains as to why we find an underrepresentation of neurons that are bottom-up excited and top-down inhibited. This is likely a consequence of the fact that we used mismatch responses, a negative prediction error, to classify the neurons. Using the unbiased measure of correlation with visual flow and locomotion (**Figure 6B**), we find a more symmetric split between the two types of responses. Testing the model proposed in the predictive processing framework will require addressing the question of whether L2/3 neurons that do not exhibit spiking responses in this paradigm, would code for a different kind of prediction error.

In sum, we show that visual and locomotion-related inputs to L2/3 neurons in visual cortex underlie visuomotor mismatch signals, and that this computation is likely specific to L2/3. In addition, we find that there are two functional types of neurons whose responses are consistent with signaling either positive or negative prediction errors. We speculate that L5/6 neurons integrate over prediction error inputs by being net inhibited by dMM neurons and net excited by hMM neurons. It is conceivable that the two functional neuron types in L2/3 are associated with different gene expression profiles. Identifying such markers would allow us to test the hypotheses put forward in the predictive processing framework.

## SUPPLEMENTARY FIGURES

**Figure S1.**
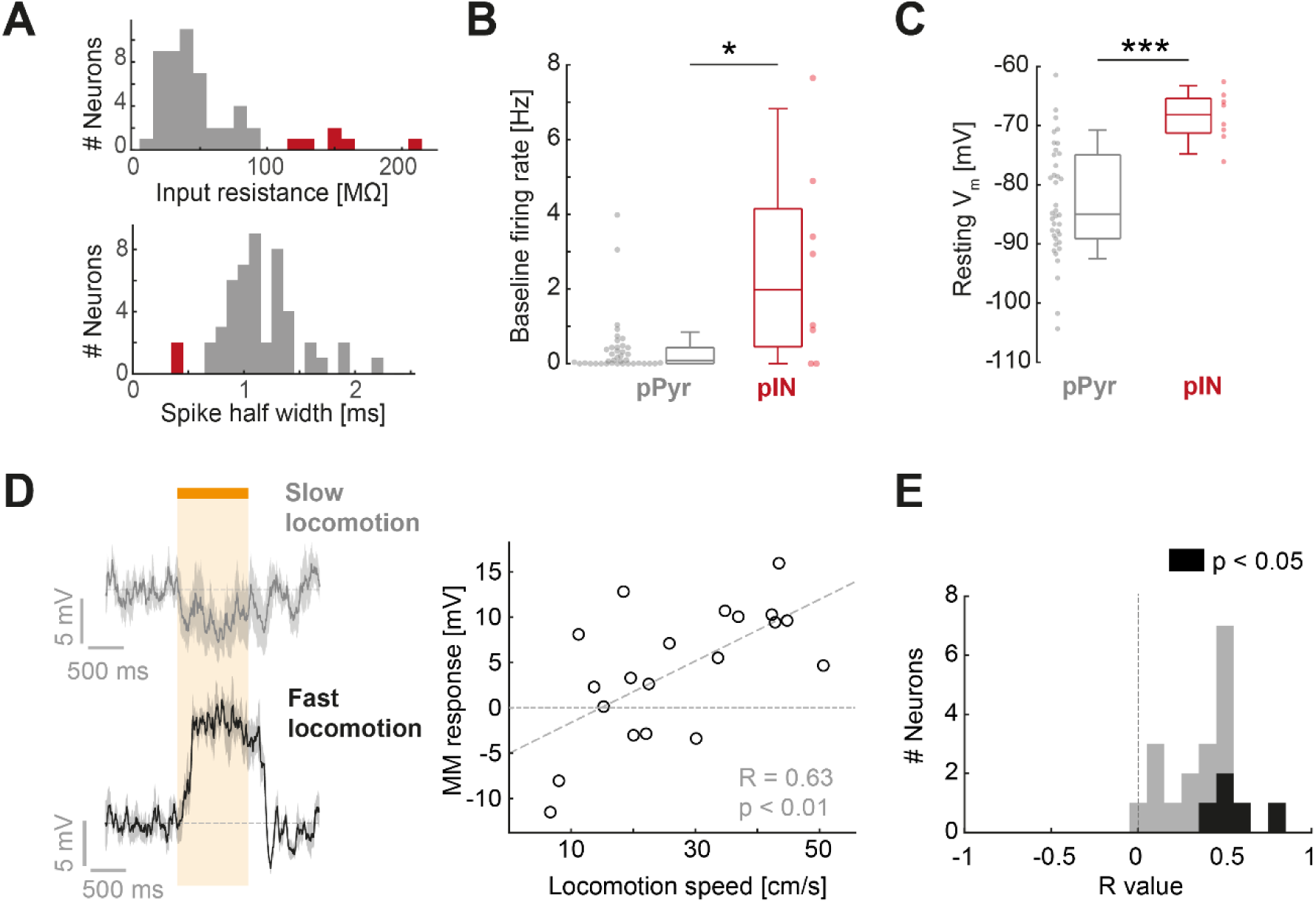
Exclusion of putative interneurons and correlations between mismatch responses and locomotion speed. Related to Figure 1. (**A**) Distribution of input resistance (top) and spike half-width (bottom) of the entire dataset, regardless of recording depth. Marked in red are neurons we excluded as potential interneurons, either based on input resistance > 100 MΩ or a spike half-width < 0.6 ms. Neurons excluded using each criterion did not overlap. Excluded neurons showed other electrophysiological properties that differed from the remaining dataset and were consistent with interneuron properties (see **B-C**). (**B**) Comparison of baseline firing rate (during stationary periods) for putative excitatory neurons versus the excluded putative interneurons. Firing rates for putative interneurons were significantly more variable (p < 10^−3^, Brown-Forsythe test), and significantly higher than in putative excitatory neurons (p < 0.03, Wilcoxon rank sum test). (**C**) As in **B**, but for resting membrane potential. Membrane potentials in putative interneurons were significantly less variable (p < 0.05 Bartlett test), and significantly more depolarized than for putative excitatory neurons (p < 10^−3^, Wilcoxon rank sum test). (**D**) Average mismatch responses from a dMM neuron at different locomotion speeds. Top: Average over 5 trials with lowest locomotion speed. Bottom: Average over 5 trials with highest locomotion speed. Shading indicates SEM. Right: Scatter plot between locomotion speed and mismatch response for the same neuron. Gray dashed line indicates the linear regression. (**E**) Histogram of R values for the correlation between locomotion speed and mismatch response in 19 neurons with more than 10 mismatch trials. In black are neurons with a significant correlation (p < 0.05).

**Figure S2.**
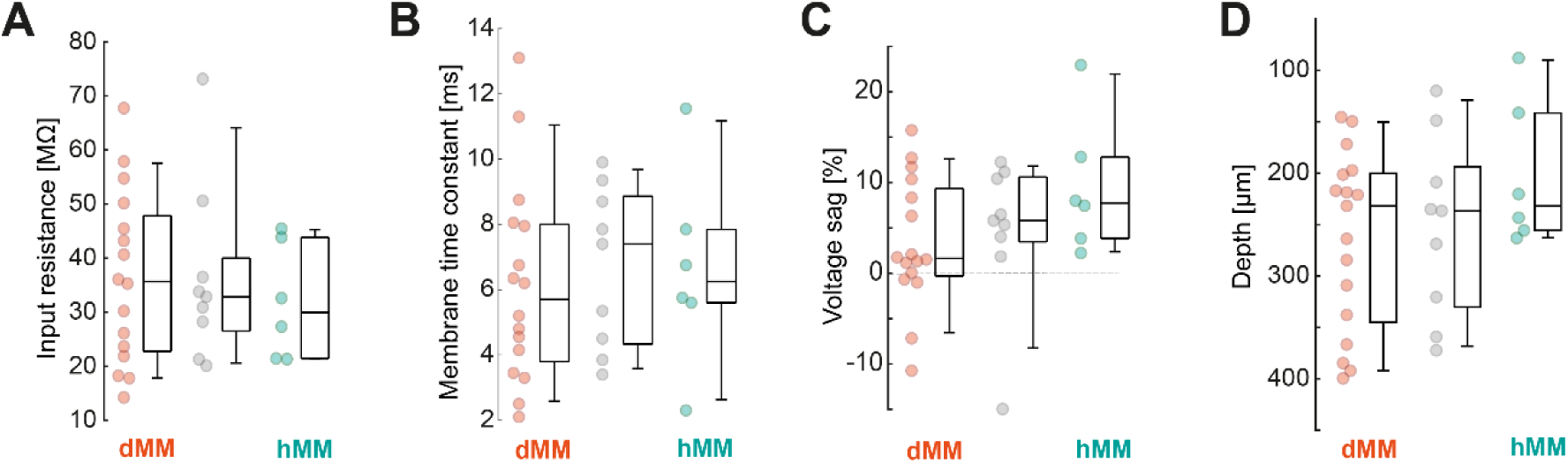
Comparison of properties between dMM and hMM neurons. Related to Figure 2. (**A**) Input resistances across the three groups of neurons. There was no significant difference between hMM and dMM groups (p = 0.54, Student’s t-test). (**B**) Membrane time constants across the three groups of neurons. There was no significant difference between hMM and dMM groups (p = 0.75, Student’s t-test). (**C**) Voltage sag during a -0.4 nA current step (a measure of I_h_ current) across the three groups of neurons. Voltage sag was larger in the hMM neurons compared to dMM neurons (likely as a consequence of the more hyperpolarized resting membrane potentials), though this did not reach significance (p = 0.09, Student’s t-test). (**D**) Vertical depth of recording from brain surface across the three groups. There was no significant difference between hMM and dMM groups (p = 0.06, Student’s t-test).

**Figure S3.**
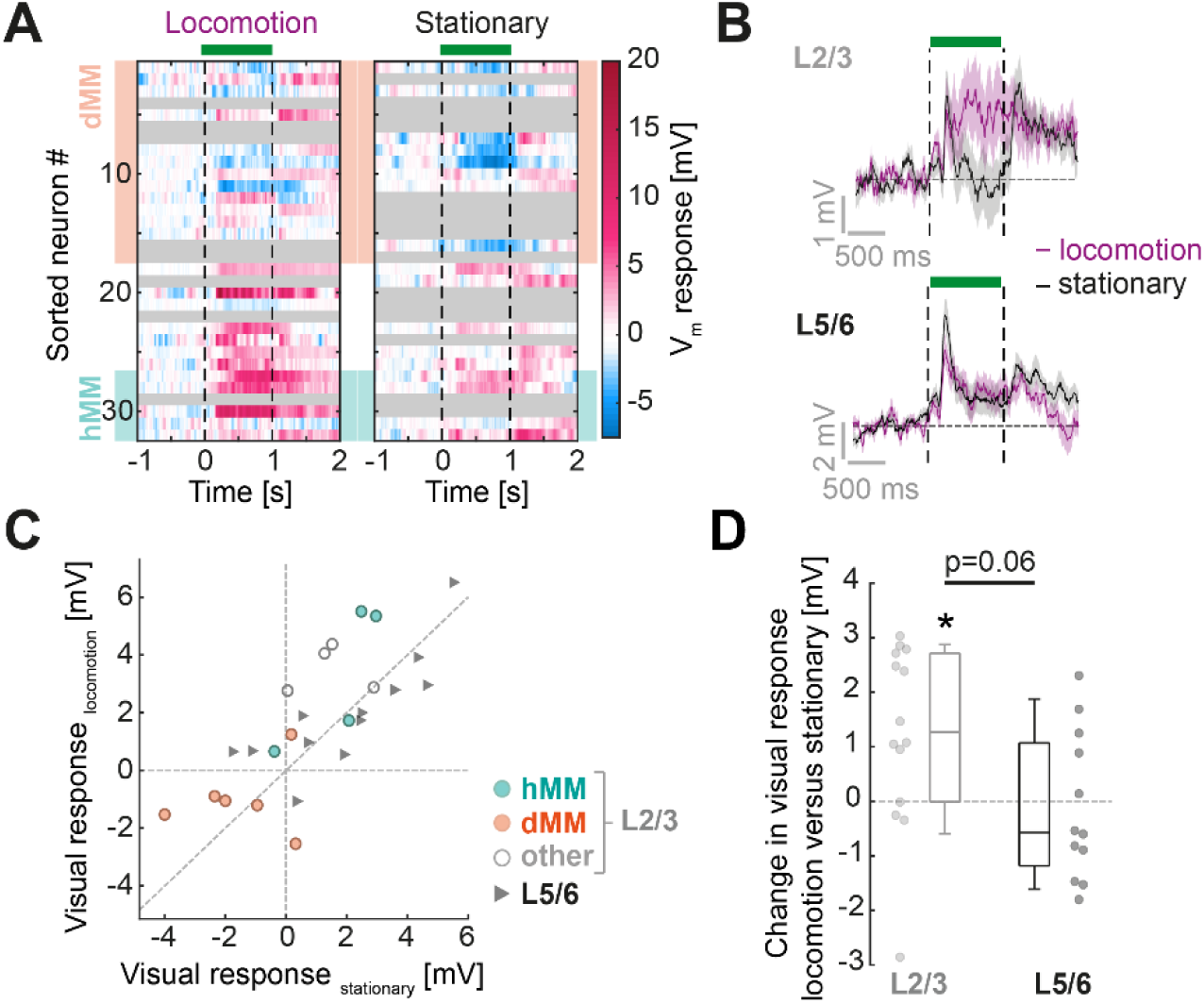
Effect of locomotion state on visual flow responses. Related to Figure 3. (**A**) Heatmaps of subthreshold visual responses of L2/3 neurons sorted by mismatch response (as in **Figure 2C**), during locomotion (left), or during stationary periods (right). Gray marks neurons for which we did not have at least five visual flow presentations. Orange shading indicates dMM neurons and turquoise shading indicates hMM neurons. (**B**) Average visual flow response of L2/3 cells (top) and L5/6 cells (bottom) during locomotion (purple) and stationary periods (black). Shading shows SEM. Only neurons with at least five trials in each category were included. (**C**) Scatter plot of visual flow responses during stationary periods and visual flow responses during locomotion periods for all neurons with at least five trials in each category. (**D**) Change in visual flow response between locomotion and stationary periods for L2/3 neurons and L5/6 neurons. L2/3 neurons showed a significantly more positive responses during locomotion compared to stationary periods (p < 0.02, paired t-test). L5/6 neurons did not show this effect (p = 0.78, paired t-test). Changes were higher for L2/3 neurons versus L5/6 neurons (p = 0.06, Student’s t-test).

**Figure S4.**
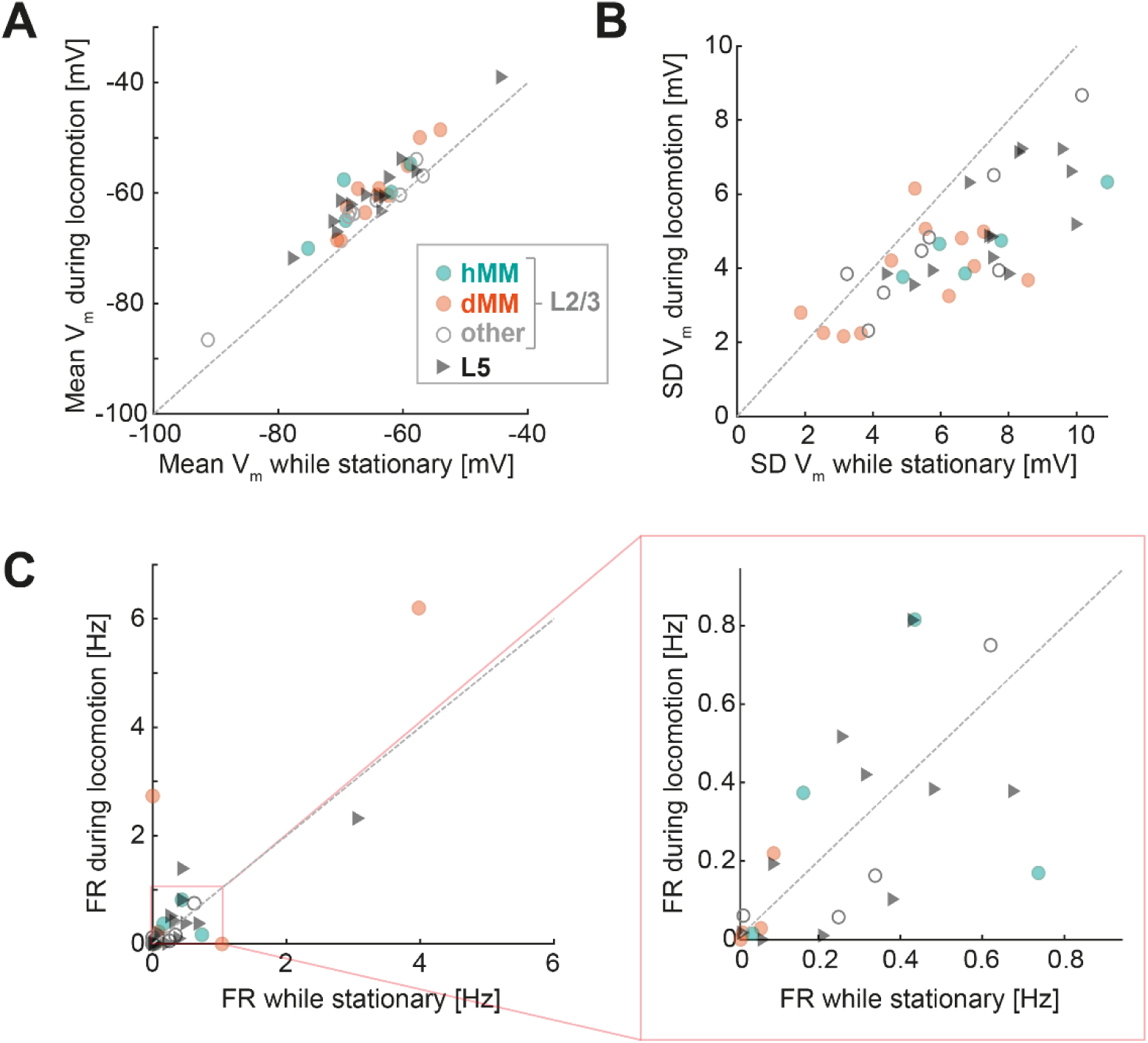
Membrane potential dynamics and firing rate changes between stationary periods and locomotion. Related to Figure 4. (**A**) Mean membrane potential (V_m_) during stationary periods versus that during locomotion. All neurons showed depolarization of membrane potential during locomotion. (**B**) As in **A**, but for the standard deviation (SD) in membrane potential. (**C**) As in **A**, but for firing rates (FR). Right plot shows an expanded version of the left.

**Figure S5.**
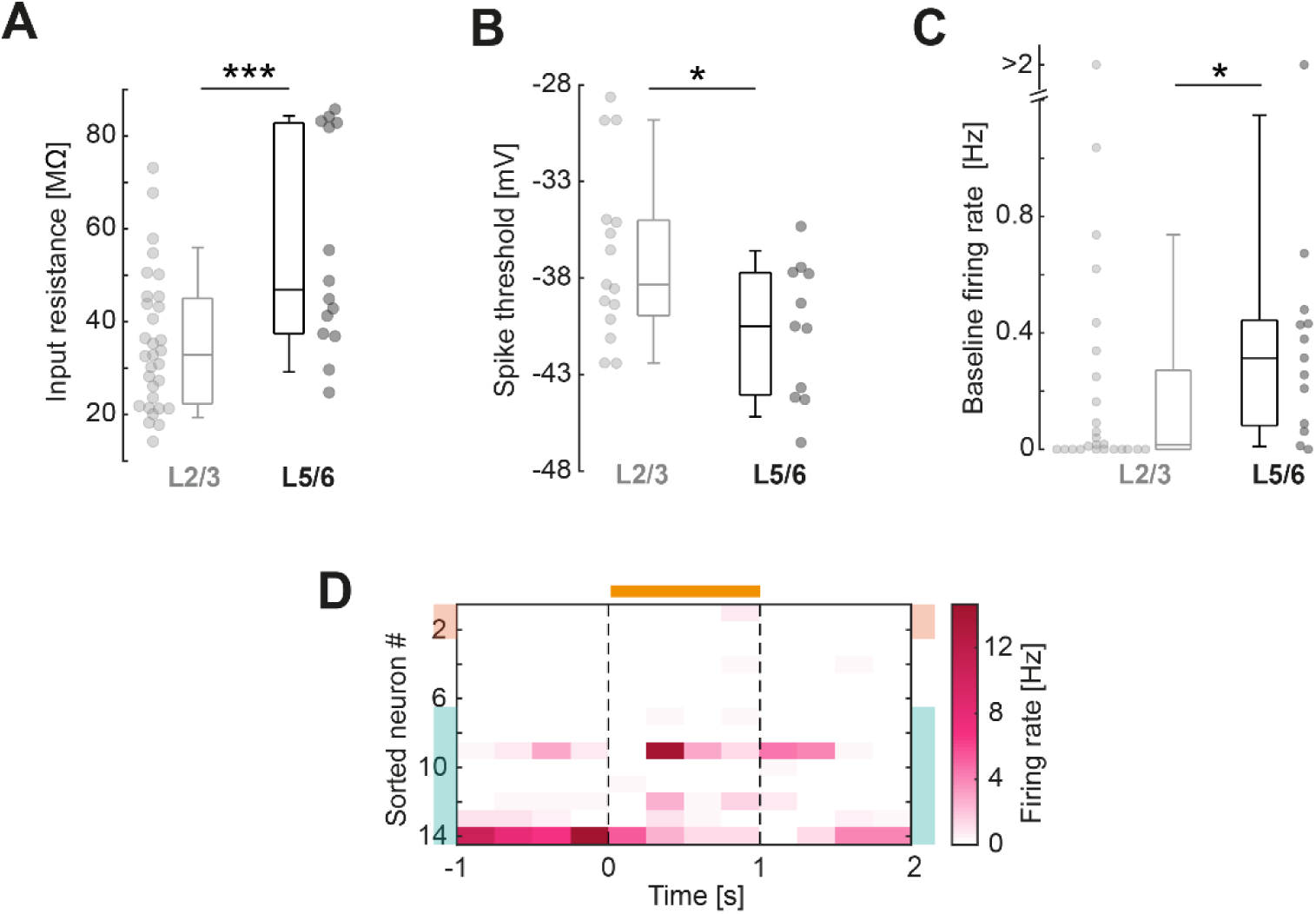
Comparison of properties between putative L5/6 and L2/3 excitatory neurons. Related to Figure 5. (**A**) Input resistance was significantly higher in L5/6 neurons than in L2/3 neurons (p < 10^−3^, Student’s t-test). (**B**) Spike threshold was significantly higher in L2/3 neurons than in L2/3 neurons (p < 0.03 Student’s t-test). (**C**) Baseline firing rate was significantly higher in L5/6 neurons than in L2/3 neurons (p < 0.03, Wilcoxon rank sum test). (**D**) Heatmap of average spike counts aligned to mismatch events for L5/6 neurons. Heatmap is sorted according to subthreshold mismatch responses, as in **Figure 4**.

**Figure S6.**
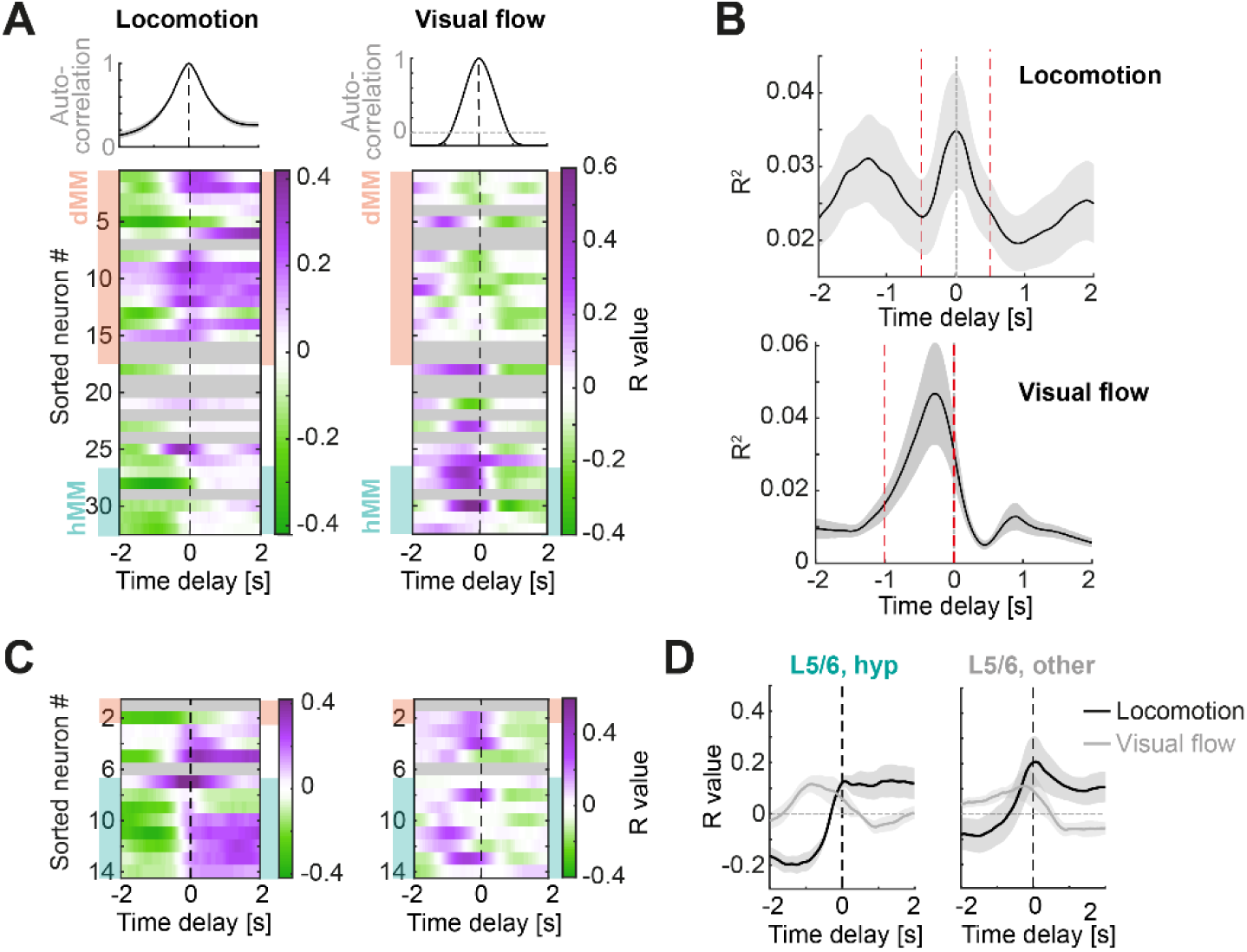
Additional data for cross correlations between visual flow, locomotion and membrane potential. Related to Figure 6. (**A**) Top plots: Average autocorrelations for locomotion (left) and visual flow (right). Heatmaps show cross-correlations for each L2/3 neuron between locomotion and membrane potential (left), and visual flow and membrane potential (right). Heatmaps are sorted by mismatch response as in main figures. All analyses excluded stationary periods. Note for all panels, negative time values indicate a lag of V_m_ relative to locomotion/visual flow. Shading indicates SEM over neurons. (**B**) Average R^2^ for all cross correlations (L5/6 and L2/3 pooled, n = 34) for locomotion and V_m_ (top) and visual flow and V_m_ (bottom). Red 1 s window indicates the time delay window used to calculate the average R value for each neuron (as plotted in **Figure 6**) – approximately centered around the peak R^2^ for locomotion and visual flow separately. (**C**) As in **A**, but for L5/6 neurons. (**D**) Average cross correlations between locomotion speed (black) or visual flow speed (gray) and membrane potential for the 7 L5/6 neurons which hyperpolarize during mismatch (‘hyp’), and the remaining 5 neurons (‘other’, including 2 depolarizing neurons) L5/6 neurons.

## METHODS

### Animals and surgery

All animal procedures were approved by and carried out in accordance with guidelines of the Veterinary Department of the Canton Basel-Stadt, Switzerland. Mice were anesthetized using a mix of fentanyl (0.05 mg/kg), medetomidine (0.5 mg/kg) and midazolam (5 mg/kg). Analgesics were applied perioperatively. Lidocaine was injected locally on the scalp (10 mg/kg s.c.) prior to surgery, while metacam (5 mg/kg, s.c.), and buprenorphine (0.1 mg/kg s.c.) were injected just after completion of the surgery. An incision was made in the skin above the cranium, and the periosteum completely removed from the skull. The surface of the skull was roughened with a dental drill. To optimize stability of the brain for later recordings, a blunt tool was used to apply force to one of the intra-parietal skull plates just anterior to bregma until small forces applied to either intra-parietal or parietal skull plates did not result in relative movement of the bones. In this position, layers of tissue glue (Histoacryl, B.Braun, Germany) were used to fuse the skull plates along the sutures. Tissue glue was then applied to the whole skull surface, and a custom-made stainless-steel head bar was glued to the skull. At this point, right V1 was marked at 2 to 3 mm lateral, just anterior to the lambdoid suture. Dental cement was used to fix the head-bar in place and build a recording chamber around V1. Anesthesia was then antagonized (Flumazenil, 0.5 mg/kg and Atipamezole, 2.5 mg/kg i.p.), and the mouse was allowed to recover for 3 days, with buprenorphine injected as before on days 1 and 2 following surgery.

### Whole cell recordings

Micropipettes (5 to 8 MΩ) were fabricated using a PC-100 puller (Narishige, Tokyo, Japan) from 1.5 mm diameter filamented borosilicate glass (BF150-86-10, Sutter, California, USA). A small 1 mm craniotomy and durectomy were made over the right primary visual cortex (2-3 mm lateral from the midline, just anterior to the lambdoid suture) under isoflurane anesthesia. To stabilize the brain, the craniotomy was then covered in a layer (0.5 - 1 mm) of 4% low-melting point agar (A9793, Sigma-Aldrich), dissolved in bath recording solution. The recording chamber was then submerged in bath recording solution (126 mM NaCl, 5 mM KCl, 10 mM HEPES, 2 mM MgSO_4_, 2 mM CaCl_2_, 12 mM glucose, brought to pH 7.4 using NaOH, with a final osmolarity 280-290 mOsm). The mouse was allowed to recover from isoflurane anesthesia for at least 20 minutes head-fixed prior to recordings, which were only attempted after the mouse had displayed regular locomotion behavior. Whole cell recordings were performed blindly by lowering the micropipette, back-filled with intracellular recording solution (135 mM KMeSO_3_, 5 mM KCl, 0.1 mM EGTA, 10 mM HEPES, 4 mM Mg-ATP, 0.5 mM Na_2_-GTP, 4 mM Na_2_-phosphocreatine, brought to pH 7.3-7.4 with KOH, with an osmolarity 284 to 288 mOsm), through the agar and 50 µm into the tissue with high pressure (>500 mbar) applied to the micropipette. Micropipette resistance was monitored in voltage clamp via observing the electrode current while applying 15 mV square pulses at 20 Hz. Brain entry was detected by a step change in the current (Margrie et al., 2002), and at this point the descent axis was zeroed. Once a depth of 50 µm from the surface was reached, pipette pressure was lowered to 20 mbar and neuron hunting began. This consisted of advancing the electrode in 2 µm steps until a substantial and progressive increase in pipette resistance was observed for at least 3 consecutive steps. Pressure in the pipette was then rapidly lowered to 0 mbar, and often a small negative pressure was applied to aid in forming a gigaohm seal. Once this was achieved, the pipette was then carefully retracted by up to 4 µm, and break-in achieved using suction pulses. Electrophysiological properties were determined in the first 60 s of the recording using a series of current steps from -0.4 to 0.3 nA, and the evoking of action potentials was used to confirm the neuronal nature of the cell. All recordings took place in current clamp mode. Recordings were terminated if series resistance displayed a substantial increase, as monitored by 25 ms current pulses between -0.1 and -0.25 nA applied at 1 Hz throughout the recording. Pipette capacitance and series resistance were not compensated. Data were acquired and Bessel low-pass filtered below 4 kHz using a MultiClamp amplifier (Molecular Devices, California USA) and digitized at 20 kHz via custom written LabView software. A junction potential of -8.5 mV was measured for our solutions, and all values reported in the manuscript have been corrected for this. On average, there was an access resistance of 58 ± 22 MΩ, and a recording duration of 14 ± 5 minutes.

### Virtual reality

During all recordings, mice were head-fixed in a virtual reality system as described previously (Leinweber et al., 2014). Briefly, mice were free to run on an air-supported polystyrene ball, the rotation of which was coupled to linear displacement in the virtual environment projected onto a toroidal screen surrounding the mouse. From the point of view of the mouse, the screen covered a visual field of approximately 240 degrees horizontally and 100 degrees vertically. The virtual environment presented on the screen was a virtual tunnel with walls consisting of continuous vertical sinusoidal gratings. Prior to the recording experiments, mice were trained in 1 to 2-hour sessions for 5 to 7 days, until they displayed regular locomotion.

### Visual stimuli

During the first segment of each recording, visual flow feedback was coupled to mouse’s locomotion speed. At random intervals averaging at 7 s, 1 s long pauses in visual feedback were presented (referred to as ‘mismatch’ stimuli). After at least 3 minutes of this protocol, the visual feedback was stopped (i.e. no visual flow coupled to locomotion speed), and instead 1 s full-field fixed-speed visual flow stimuli were presented at random intervals (mean ± SD, 8.1 ± 1.3 s), regardless of locomotion behavior. In a subset of recordings (12 of 27), these stimuli all had one fixed visual flow speed, and in the remaining subset (15 of 27), four different visual flow speeds were presented in a pseudorandom sequence.

### Data Analysis

All data analysis was performed using custom written Matlab 2019a (Mathworks) code.

### Comparison statistics

For each unpaired two-sample comparison, first a Lilliefors test was used to test whether the distribution was normal. If so, a Bartlett test was used to determine whether the two samples had an equal variance. If both conditions were satisfied (p > 0.05), a Student’s t-test was used to determine whether there was a significant difference between the two groups. If either condition was violated, a Wilcoxon rank-sum test was used instead. To test for significant differences in variance in non-normally distributed samples, a Brown-Forsythe test was performed. Box and whisker plots are all drawn such that the box represents the inter-quartile range and median, and the whiskers represent the 10^th^ and 90^th^ percentiles. For correlations, as in **Figures 3E, 4D** and **6A**, a method of fitting robust to outliers was used (Matlab function Fitlm) based on bisquare weighting of residuals.

### Cell numbers

In total, we recorded from 54 neurons. Of these, 6 neurons were excluded as putative interneurons, as they had an input resistance higher than 100 MΩ. A subset of interneuron types (e.g. somatostatin-expressing neurons) have been described to have high input resistances. Two neurons were excluded due to spike half-widths below 0.6 ms (putative parvalbumin-expressing neurons). Consistent with the excluded neurons being interneurons, other electrophysiological features differed between these excluded neurons and the remaining putative excitatory neurons, including higher baseline spike rates, more depolarized resting membrane potential and lower spike thresholds (**Figure S1**). 32 of the remaining 46 putative excitatory neurons were recorded at a vertical depth of less than 400 µm below the brain surface. We refer to these as putative L2/3 neurons. Of these, 28 underwent both the visuomotor coupled and open-loop parts of the protocol, and the remaining 4 underwent only the former part. Of the 28 neurons with both parts of the protocol, 16 were presented with visual stimuli of four different speeds, while the remaining 12 were presented with only one visual flow speed. The neurons for which we do not have data in the open-loop condition are represented as uniform gray on response heatmaps of visual flow and locomotion onset response. 14 neurons were recorded at vertical depths lower than 480 µm, with a maximum depth of 723 µm. We refer to these as putative L5/6 neurons. Of these, 13 underwent both the visuomotor coupled and open-loop parts of the protocol, and the remaining 1 underwent only the former part. Of the 13 neurons with both parts of the protocol, 12 were presented with four different visual flow stimuli of different speeds, and the remaining 1 was presented visual flow stimuli of one speed only.

### Electrophysiological properties

A series of current steps from -0.4 to 0.3 nA were applied to the neuron at least 3 times at the beginning of the recording to determine input resistance. The total resistance was calculated by averaging the voltage response for each current step value and measuring the slope between the average response 25-125 ms after current step onset against the injected current. This was done separately for negative and positive current injection, as the former consistently showed a lower resistance than the latter (in part due to voltage sag during negative current injection). Next, access resistance was calculated by taking the slope between the current injected and the voltage response in the first 1 ms. Input resistance was calculated as the difference between total resistance and access resistance. Resting membrane potential was defined where the current-voltage slope crossed the voltage-axis (at zero current). Note that resting membrane potential would often depolarize by a few mV before stabilizing during the first 3-5 minutes of the recording, presumably as the intracellular solution diffuses throughout the neuron. As such, membrane voltages read out at later time points (e.g. in **Figure S4**) are different to the resting membrane potential assessed just after break-in. Voltage sag was calculated as the difference in the average voltage response to a -0.4 nA step 15 to 25 ms after current step onset and 150 to 250 ms after onset. Note that holding voltage was not controlled, so resting membrane potential differences account for some of the variance (R^2^ = 0.24) in the voltage sag measurement. The membrane time constant was estimated by finding the time at which the voltage change first exceeded 1-1/e of the difference between 1 ms after current step (−0.1 nA) onset and steady state (estimated as the voltage in the window 45-50 ms after the onset).

### Spike properties and subtraction

Spikes were detected from peaks exceeding -30 mV during zero current application from across the entire recording. Spikes with an amplitude less than 30 mV were rejected. The remaining spikes were then averaged together to get an average spike waveform. The spike threshold was determined as the voltage value of the average spike waveform at the time of the peak rate of change of the slope of the spike waveform. Spike amplitude was measured as the difference between value of the peak of the spike and that of the threshold. For spike threshold analysis, only naturally occurring spikes were included in the analysis, and not the spikes evoked during current injection. Thus, for silent neurons these values are missing. Spike half-width was then measured as the duration the average spike waveform exceeded half of the spike amplitude. For all average membrane potential response plots, spikes were removed from membrane potential traces by replacing them with a linear interpolation from the membrane potential recorded 2 ms prior to spike peak, and that recorded 3 ms after spike peak. Voltage responses to the 25 ms current pulses used to track access resistance were removed similarly, by replacing them with a linear interpolation from the time just before the current pulse turned on to that 70 ms later.

### Mismatch responses

Presentation of visual flow halts were independent of locomotion speed. As only halts during non-zero visual flow speed would result in a change, mismatch events were defined as visual flow halts that occurred during an average locomotion speed exceeding 4 cm/s in the 1 s prior to and during mismatch stimulus. To calculate average V_m_ responses, spikes and current pulses were subtracted as described above and the average voltage in a window 1 s prior to mismatch was subtracted from the V_m_ for each trial. The mean of all resulting V_m_ traces was then taken to generate the average response. The average response for each neuron was taken as the mean V_m_ response during the entire 1 s of mismatch presentation. The significance of this response was determined by performing a paired Student’s t-test between the average V_m_ 1 s before mismatch, and that in the 1 s during mismatch for all included trials. Spiking responses were computed based on the same mismatch events and generated by taking the mean spike count in 250 ms time bins aligned to mismatch onset.

### Locomotion dynamics and correlations with V_m_

To measure membrane potential average and variance, as well as spiking activity during locomotion and stationary periods (**Figure S4**), only the data from the open-loop condition was used. The data was binned into 500 ms time bins, and the spike count, median membrane potential and locomotion speed were calculated for each time bin. The locomotion speed trace was smoothed in a 1 s time window prior to this calculation. For quantification of V_m_ during stationary periods, all V_m_ values corresponding to times when the locomotion speed was below a threshold of 4 cm/s were pooled, and the mean and standard deviation of these values were calculated for each neuron. The same was then done for locomotion periods where the locomotion speed exceeded the 4 cm/s threshold.

### *Calculation of cross correlations* (Figures 4E, 4F and S6)

Cross correlations were calculated between membrane potential and locomotion for the open-loop condition only. For this, the locomotion trace and visual flow trace recorded at 1 kHz were smoothed using a 500 ms time window. Membrane potential was binned in 1 ms time windows. Times when the mouse was stationary (locomotion < 4 cm/s), and times where visual flow stimuli were presented (as well as 1 s after the presentation) were excluded. We then computed the cross-correlation between locomotion trace and membrane potential in a window of -2000 to +2000 ms. A similar procedure was used for the cross correlation between membrane potential and visual flow, again excluding periods when the mouse was stationary. For each neuron, the overall correlation coefficient for the Locomotion-V_m_ correlation was taken as the average correlation for time delays between -500 and +500 ms, as this is where the cross correlation averaged across the L2/3 and L5/6 samples combined peaked (**Figure S6B**). For each neuron, the overall correlation coefficient for the visual flow-V_m_ correlation was taken as the average correlation for time delays between -1000 and 0 ms, as this is where the cross correlation averaged across the L2/3 and L5/6 samples combined peaked (**Figure S6B**). Only neurons with at least 25 s of locomotion in absence of visual flow were included in these analyses (n = 22 L2/3, n = 12 L5/6).

To compare correlations for L5/6 and L2/3 datasets (**Figure 6**), we calculated an interaction angle for each neuron as the arcus tangent of the ratio of the locomotion-V_m_ correlation to the visual flow-V_m_ correlation. Polar histograms were then made for the L2/3 and L5/6 datasets separately (**Figure 6B**). The neuron counts for each time bin were then correlated between L5/6 and L2/3 datasets, generating an R value of -0.11. To test if this anticorrelation was significant, L5/6 and L2/3 angles were pooled, and random subsets corresponding to the sample sizes of the L5/6 and L2/3 datasets were drawn out. An R value was then calculated for the correlation between the resulting two polar histograms. This was repeated 10000 times to generate a distribution of R values. Only 0.6 % of the distribution had a negative R value below -0.11.

To determine how well the correlations for visual flow speed and locomotion speed predicted a neuron’s mismatch response (**Figure 6C**), we computed the correlation between the difference of the two R values (R value_locomotion_ - R value_visual_) and the mismatch responses, for L2/3 and L5/6 neurons separately. As a shuffle control, we then randomly permuted the visual flow correlation and locomotion correlation values across neurons 100000 times to create a shuffle distribution.

### Visual responses

Visual onsets were defined as e the visual flow trace crossing a threshold of 0.8 cm/s. For average visual responses, the membrane potential for each presentation was baseline subtracted by the average V_m_ in the 1 s prior to visual flow onset. The response was then averaged across all trials, regardless of locomotion behavior. For the subset of neurons which were shown four different visual flow speeds, responses were averaged regardless of visual flow speed. To determine the effect of locomotion on visual flow responses (**Figure S3**), visual flow presentations were separated according to locomotion speed: locomotion trials were defined as trials in which the average locomotion speed in the 1s prior and 1 s during visual flow both exceeded 4 cm/s. Stationary trials were defined as trials in which the locomotion speed in these two epochs both were less than 4 cm/s. For correlations between visual flow speed and visual response, only trials in which the mouse was locomoting above 0.8 cm/s were included, and an R value for the correlation between the visual flow speed and membrane potential response across trials was generated for each neuron.

### Correlation between visual flow response and mismatch response

For the correlation between mismatch response and visual response across cells (**Figure 3E**), we first calculated the correlation between the mismatch response (as averaged across the 1 s of mismatch) versus the visual response (as averaged across the 1 s of visual response). To account for any response to visual flow offset, we took the visual offset response as the average membrane potential response in the 1 s after visual flow offset, normalized to the average membrane potential in the 1 s prior to visual flow onset. This visual flow offset response was then subtracted from the mismatch response, and the correlation was calculated between the resulting values and the visual flow response.

### Locomotion onset responses

Locomotion velocity was first smoothed using a 1 s time window. To determine the time of locomotion onsets, we detected where the smoothed locomotion velocity crossed a threshold of 0.8 cm/s. We then excluded any onsets where the smoothed locomotion velocity 1 s prior exceeded 2.5 cm/s, and the velocity 1 s after onset was less than 2.5 cm/s. Average locomotion onset responses were then calculated for each neuron where there were at least 2 locomotion onsets. These were normalized by subtracting the average membrane potential in the 2.5 s prior to locomotion onset for each trial before averaging all traces. Locomotion onset responses were taken for each neuron as the average response 0-6 s after locomotion onset.

## ACKNOWLEDGEMENTS

We thank Rainer Friedrich, Mahesh Karnani, Jesse Jackson, Loreen Hertäg and Andreas Keller for helpful discussion and comments on earlier versions of this manuscript. We thank Andreas Lüthi for lending us research equipment, and all the members of the Keller lab for discussion and support. This project has received funding from the Swiss National Science Foundation (GBK), the Novartis Research Foundation (GBK&RJ), the Human Frontier Science Program (RJ), and the European Research Council (ERC) under the European Union’s Horizon 2020 research and innovation programme (grant agreement No 865617) (GBK).

## AUTHOR CONTRIBUTIONS

RJ designed and performed the experiments and analyzed the data. Both authors wrote the manuscript.

